# Human olfactory neuronal cells through nasal biopsy: molecular characterization and utility in brain science

**DOI:** 10.1101/2022.09.23.509290

**Authors:** Kun Yang, Koko Ishizuka, Andrew P. Lane, Zui Narita, Arisa Hayashida, Yukiko Y. Lema, Emma Heffron, Haydn Loudd, Maeve Schumacher, Shin-Ichi Kano, Toshifumi Tomoda, Atsushi Kamiya, Minghong Ma, Donald Geman, Laurent Younes, Akira Sawa

## Abstract

Biopsy is crucial in clinical medicine to obtain tissues and cells that directly reflect the pathological changes of each disease. However, the brain is an exception due to ethical and practical challenges. Nasal biopsy, which captures the olfactory neuronal epithelium, has been considered as an alternative method of obtaining neuronal cells from living patients. Multiple groups have enriched olfactory neuronal cells (ONCs) from biopsied nasal tissue. ONCs can be obtained from repeated biopsies in a longitudinal study, providing mechanistic insight associated with dynamic changes along the disease trajectory and treatment response. Nevertheless, molecular characterization of biopsied nasal cells/tissue has been insufficient. Taking advantage of recent advances in next-generation sequencing technologies at the single-cell resolution and related rich public databases, we aimed to define the neuronal characteristics, homogeneity, and utility of ONCs. We applied single-cell and bulk RNA sequencing for ONCs, analyzing and comparing the data with multiple public datasets. We observed that the molecular signatures of ONCs are similar to those of neurons, distinct from major glial cells. The signatures of ONCs resemble those of developing neurons and share features of excitatory neurons in the prefrontal and cingulate cortex. The high homogeneity of ONCs is advantageous in pharmacological, functional, and protein studies. Accordingly, we provide two proof-of-concept examples for functional and protein studies, solidifying the utility of ONCs in studying objective biomarkers and molecular mechanisms for brain disorders. The ONCs may also be useful in the studies for the olfactory epithelium impairment and the resultant mental dysfunction elicited by SARS-CoV-2.

**SIGNIFICANCE STATEMENT:** To study dynamic changes and underlying mechanisms along disease trajectory and treatment response in neuropsychiatric disorders, olfactory neuronal cells (ONCs) enriched from biopsied nasal tissue may provide a crucial tool. Because ONCs can be obtained from repeated biopsies in a longitudinal study, this tool has been believed to be useful and complementary to postmortem brains and induced pluripotent stem cell-derived neurons. Nevertheless, molecular characterization of biopsied nasal cells/tissue has been insufficient, which hampers a broader use of this resource. Taking advantage of recent advances in next-generation sequencing technologies at the single-cell resolution and related rich public databases, the present study defines ONCs’ neuronal characteristics, homogeneity, and unique utility for the first time.

## INTRODUCTION

Biopsy is an essential procedure of clinical medicine where we obtain tissues and cells that are supposed to directly reflect pathological changes of each disease. This procedure allows us to analyze pathological deficits at the molecular and cellular levels, contributing to precise diagnosis and proper choice of medications (Joyce et al., 2012; Elston et al., 2016; Schiffer, 2018). These biospecimens are also useful in drug discovery (Basik et al., 2013; Jafari et al., 2014; Morrison, 2017; Siravegna et al., 2017; Fernández-Lázaro et al., 2020). By using such biospecimens in combination with frontline multi-omics, such as genomic, epigenomic, and proteomic approaches, many successful basic and translational research projects have expanded scientific knowledge of disease pathophysiology. The successful precedents include the Encyclopedia of DNA Elements (ENCODE) (Consortium, 2004) and the Cancer Genome Atlas (TCGA) (Tomczak et al., 2015).

Nevertheless, because of the challenges of brain biopsy at both practical and ethical levels, this approach to brain disorders is not straightforward. To overcome this limitation, a translational application of induced pluripotent stem cells (iPSCs) has been considered. iPSC-derived neurons have been successfully used in many basic studies because these neurons are expected to maintain cell-type-specific molecular signatures. A possible drawback of the iPSC-based methodology is that reprogramming in iPSC protocols could affect molecular signatures associated with disease states, limiting the utility of iPSCs in longitudinal studies (De Los Angeles et al., 2021). In addition, the scalability of iPSCs faces technical and budget challenges (Huang et al., 2019; Ortuño-Costela et al., 2019). Furthermore, many iPSC protocols lead to heterogeneous sets of neurons and non-neuronal cells in culture, which hampers biochemical and pharmacological approaches that need homogeneous cell populations, although single-cell assays may complement this limitation to some extent.

Since the 1980s, the olfactory neural epithelium has also been considered as a potentially useful biospecimen for brain studies (Lovell et al., 1982; Graziadei and Monti Graziadei, 1985; Talamo et al., 1989). This epithelium, directly accessible through the nasal cavity, contains olfactory neurons and their progenitors that regenerate mature neurons throughout life (Graziadei and Graziadei, 1979). Owing to the difficulty of capturing only neural epithelium and the potential contamination of mucosal tissues in nasal biopsy, multiple groups have developed protocols to enrich olfactory neuronal cells (ONCs) from the biopsied tissue, expecting that these cells have neuronal characteristics (Mackay-Sim, 2012; Borgmann-Winter et al., 2015; Kumar et al., 2018; Rhie et al., 2018; Evgrafov et al., 2020). Our group has also published many reports using ONCs (Sawa and Cascella, 2009; Horiuchi et al., 2013; Kano et al., 2013; Mor et al., 2013; Lavoie et al., 2017; Sumitomo et al., 2018; Takayanagi et al., 2021; Namkung et al., 2023; Jaaro-Peled et al., 2022). Nevertheless, insufficient characterization of biopsied cells/tissue has limited the broader use of nasal tissues as surrogate resources for brain science.

We aimed to overcome this challenge by taking advantage of recent advances in next-generation sequencing technologies at the single-cell resolution and related rich public databases. The goals of the present study are to define the neuronal characteristics, homogeneity, and unique utility of ONCs. To address the first two questions, we conducted single-cell and bulk RNA sequencing (RNA-seq), and analyzed and compared the data with multiple public datasets.

For the third question, we demonstrate the utility of ONCs in exploring protein and functional/mechanistic signatures directly associated with clinical features. Treatment resistance (TR) is a major clinical problem in psychotic disorders, such as schizophrenia. We compared the neuronal signatures between TR and non-TR patients, demonstrating that TR may be associated with a signaling cascade of the α-amino-3-hydroxy-5-methyl-4-isoxazolepropionic acid (AMPA)*-*type ionotropic glutamate receptor. In addition to this proof-of-concept evidence, we also show another example of ONCs’ utility in demonstrating an aberrant P62 protein expression associated with aging in psychotic patients. Altogether, we propose the utility of ONCs as a neuronal resource obtained from living patients to directly study clinical feature-associated neuronal signatures and biological mechanisms.

## MATERIALS AND METHODS

### Nasal biopsy

Nasal biopsy was performed under endoscopic control with local anesthesia to the nasal cavity provided by lidocaine liquid 4% and oxymetazoline HCl 0.05% spray, followed by injection of 1% lidocaine and 1/100,000 epinephrine. The olfactory epithelium (OE) is more likely to be found further back in the nasal cavity on the septum as well as on the turbinates (Féron et al., 1998). More specifically, the probability of obtaining OE was reportedly 40 %, 76 %, 30-50 %, 73 %, and 58 % at the dorsomedial and the dorsoposterior areas of the septum, the dorsomedial and dorsoposterior areas of the superior turbinate, and the middle turbinate, respectively (Féron et al., 1998). Based on this information, three tissue pieces were taken from the OE regions of the septum from each nostril (6 pieces total) by using ethmoid forceps to avoid trauma to the cribriform plate. When accessing the septum was difficult, the superior turbinate was targeted. After the biopsy, participants were observed in the clinic for 15 to 30 min. Since this procedure was approved by Johns Hopkins IRB in 2004, we have performed over 400 biopsies without major complications.

### ONCs

**Fig. 1** summarizes the procedure for enriching ONCs from nasal biopsy. First, biopsied tissues were incubated with 2.4 U/mL Dispase II for 45 min at 37°C, and mechanically minced into small pieces. Then, the tissue pieces were further treated with 0.25 mg/mL collagenase A for 10 min at 37°C. Dissociated cells were gently suspended, and centrifuged to obtain pellets. Cell pellets were resuspended in DMEM/F12 supplemented with 10% FBS and antibiotics (D-MEM/F12 medium), and tissue debris was removed by transferring only the cell suspension into a new tube. Cells from the suspension were then plated on 6-well plates in fresh D-MEM/F12 medium (Day 0 plate). Cells floating were collected on day 2 (Day 2 plate) and further incubated on 6-well plates. On day 7, the cells floating in the Day 2 plate were collected and further incubated on 6-well plates (Day 7 plate). Through these processes, connective tissue-origin cells and non-neuronal cells that are attached to Day 0 and Day 2 plates are removed. Accordingly, in the present study, we examined cells in Day 7 plates called ONCs under the hypothesis that these are neuronal and homogeneous. ONCs were aliquoted and stored in liquid nitrogen tanks for further experiments, at which point ONCs were recovered and cultured in the D-MEM/F12 medium with a media change every 2-3 days.

**Figure 1.**
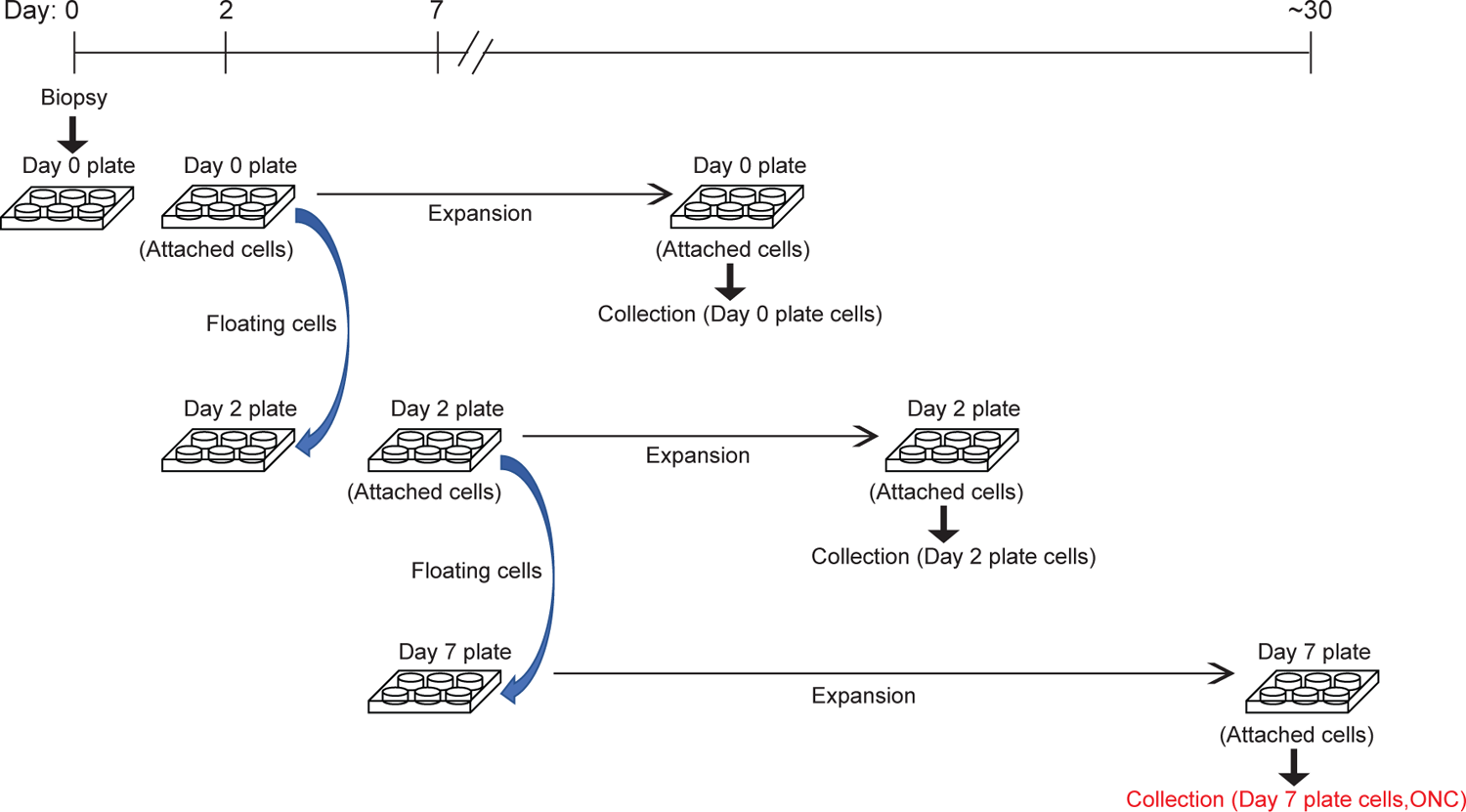
Experimental procedure for enriching ONCs from nasal biopsy. Olfactory cells are dissociated from biopsied tissues through mechanical and chemical approaches and incubated in plates. On the 2nd and 7th day of incubation, cells floating or loosely attached to the plate are transferred to new plates and further incubated until they reach confluency. These cells are collected and used as ONCs.

### Human subjects who contributed to the nasal biopsy

The present study used clinical data and biospecimens collected from study participants recruited by clinical cohorts established at the Johns Hopkins Schizophrenia Center (Kano et al., 2013; Mor et al., 2013; Kamath et al., 2018, 2019; Posporelis et al., 2018; Faria et al., 2019, 2021; Nucifora et al., 2019; Wang et al., 2019, 2023; Avigdor et al., 2021; Coughlin et al., 2021; Narita et al., 2021; Yang et al., 2021, 2022b; Etyemez et al., 2022b, 2022a). This study was conducted with the approval of the Johns Hopkins Medicine Institutional Review Boards. All study participants provided written informed consent. All study participants have no history of traumatic brain injury, cancer, abnormal bleeding, or other major physical conditions, no substance or alcohol abuse in the past three years, and no illicit drug use in the past two months. Healthy controls must have no personal or immediate family history of psychosis and do not take psychotropic medication at enrollment. Patients received the Structured Clinical Interview for DSM-IV (SCID-IV) by study team psychiatrists. Patients with chronic schizophrenia or first episode psychosis (disease onset within 24 months of initial study enrollment) were used in this study. Collateral information for patients was also gathered from available medical records.

To stratify patients with psychosis into TR group and non-TR group, we employed the criteria (the use of clozapine or no response to two or more non-clozapine antipsychotics) frequently used in many publications, including our past publications (Kayo et al., 2012; Lally et al., 2016; Wimberley et al., 2016; Howes et al., 2017; Nucifora et al., 2018; Yang et al., 2022b).

Experienced psychiatrists in the Johns Hopkins Schizophrenia Center reviewed all the medication records and clinical interview notes. Patients who switched antipsychotics due to adverse effects, poor adherence, or other reasons independent of their medication response were not classified as TR.

### External datasets as references for the homogeneity of ONCs

1. HEK293T and Jurkat cells: single-cell RNA sequencing (scRNA-seq: 10x sequencing) was conducted in cells from American Type Culture Collection (ATCC) and cultured according to ATCC guidelines (Zheng et al., 2017).
2. iPSC-derived neural mixed culture: scRNA-seq/10x sequencing was conducted in neural culture from iPSCs by a chemical induction (Earley et al., 2021).
3. iPSC-derived dopaminergic neurons: scRNA-seq/10x sequencing was conducted in fluorescently activated cell sorting (FACS)-purified tyrosine hydroxylase (TH) neurons differentiated from iPSCs (Earley et al., 2021).
4. iPSC-derived sensory neurons: scRNA-seq/10x sequencing was conducted in nociceptive sensory neurons induced from iPSCs (Plumbly et al., 2022).
5. iPSC-derived motor neurons: scRNA-seq/10x sequencing was conducted in spinal motor neurons induced from iPSCs (Thiry et al., 2020).
6. Postmortem subgenual anterior cingulate cortex (sACC) and dorsolateral prefrontal cortex (DLPFC): single-nucleus RNA sequencing (snRNA-seq: 10x sequencing) was conducted in tissues from healthy subjects without neurological disorders (Tran et al., 2021).

### External datasets as references for the neuronal characteristics of ONCs

i. Postmortem sACC, DLPFC, hippocampus, nucleus accumbens, and amygdala: snRNA-seq/10x sequencing was conducted in the tissues of healthy subjects without neurological disorders (Tran et al., 2021).
ii. Postmortem prefrontal cortex: snRNA-seq/10x sequencing was conducted in the tissues of healthy subjects without neurological disorders (Velmeshev et al., 2019).
iii. Postmortem middle temporal gyrus: snRNA-seq/10x sequencing was conducted in the tissues of healthy subjects without neurological disorders (Hodge et al., 2019).
iv. Olfactory tissue: scRNA-seq/10x sequencing was conducted in fresh tissue samples from subjects undergoing endoscopic nasal surgery (Durante et al., 2020).
v. Postmortem brains at different developmental stages: bulk RNA-seq was conducted in the postmortem brains at the following developmental stages: early fetus (0-12 post-conception weeks), middle fetus (13-18 post-conception weeks), late fetus (19-40 post-conception weeks), infancy (0-2 years old), childhood (3-10 years old), adolescence (11-19 years old), and adulthood (20-40 years old), in the BrainSpan project (Miller et al., 2014).

### External datasets used as quality controls for *Seurat* cell type prediction

#### Positive controls: samples with known brain cell types

I. Brain neurons: snRNA-seq / HiSeq sequencing was conducted in the neuronal nuclei [enriched by using neuronal nuclear antigen (NeuN)] from postmortem brain tissues of a normal subject (Lake et al., 2016).
II. iPSC-derived dopaminergic neurons: scRNA-seq/10x sequencing was conducted in FACS-purified TH neurons programmed from iPSCs (Earley et al., 2021).
III. Brain microglia: scRNA-seq/10x sequencing was conducted in FACS-purified microglia from postmortem brain tissues (Gerrits et al., 2020). *Negative control: sample with known cell types not from the brain*
IV. Human peripheral blood mononuclear cells (PBMCs): scRNA-seq/10x sequencing was conducted in fresh PBMCs purchased from ALLCELLS (Villani et al., 2017).

### Bulk RNA-seq

#### Library preparation and sequencing

total RNA was isolated from the ONCs using the RNeasy Plus Mini Kit (Qiagen). RNA quality was assessed via Agilent Fragment Analyzer analysis using a High Sensitivity RNA analysis kit (DNF-472) and quantified using a Qubit 4 RNA BR kit (Thermo Fisher). RNA libraries were prepared with 500 ng total RNA. Library generation was accomplished using NEBNext Ultra II Directional RNA Library Prep Kit for Illumina (E7760 and E7490) following the NEBNext Poly(A) mRNA Magnetic Isolation Module protocol.

Libraries were enriched using 11 cycles of PCR amplification. Library quality and quantification were assessed via Fragment Analyzer analysis using a High Sensitivity NGS Kit (DNF-474) and Qubit 4. Samples were then normalized to 4 nM and pooled in equimolar amounts. Paired-End Sequencing was performed using Illumina’s NovaSeq6000 S4 200-cycle kit.

#### Data preprocessing

Sequencing reads were filtered to remove adapters, primers, and reads with low quality (base call accuracy < 90%) or shorter than 20 nt using *FastQC* (Andrews, 2010) and *cutadapt* (Martin, 2011), then mapped to the human genome version GRCh38 (Genome Reference Consortium Human Build 38) using *Hisat2* (Pertea et al., 2016). Count table was created after transcript assembly and abundance estimation using *Stringtie* (Pertea et al., 2016), then the batch effect was adjusted using *Combat* (Leek et al., 2012). Lowly expressed genes (gene counts were less than 10 in more than 30% of samples) were removed. Principal component analysis (PCA) was conducted to check outliers (one outlier was identified and removed from downstream analyses).

#### Downstream analysis

The processed bulk RNA-seq data from healthy controls were used to study the neuronal characteristics of ONCs via PCA. The processed bulk RNA-seq data from psychotic patients were used to study molecular signatures associated with TR via differential expression analysis: *DESeq2* (Love et al., 2014 p.2) with covariates including age, sex, race, tobacco usage status (Yes/No), antipsychotic dose (chlorpromazine equivalent dose estimated by the Defined Daily Doses method (Leucht et al., 2016)), duration of illness, and unknown confounders identified by surrogate variable analysis (SVA) (Leek et al., 2012). Chi-squared test was conducted to evaluate the enrichment of risk genes for psychotic disorders identified by genome-wide association study (GWAS): genes with single-nucleotide polymorphisms (SNPs) that had significant (p-value < 1.00E-08) correlations with schizophrenia [experimental factor ontology (EFO) id: MONDO_0005090] or psychosis (EFO id: EFO_0005407) in GWAS catalog database (MacArthur et al., 2017). Gene set enrichment analysis (GSEA) (Subramanian et al., 2005) and the Reactome Pathway Database (Fabregat et al., 2018) were used to identify molecular pathways altered in TR patients compared with non-TR patients. The Benjamini-Hochberg procedure was conducted for multiple comparison correction for all analyses involving multiple tests. Results with an adjusted p-value smaller than 0.05 were considered significant.

### scRNA-seq

#### Library preparation, sequencing, and data preprocessing

10xGenomic single cell 3′ library was prepared as follows: single cells were sorted into 1.5 mL tubes (Eppendorf) and counted under the microscope. Cells were loaded at 10,000-100,000 cells/chip position using the Chromium Single cell 3′ Library, Gel Bead & Multiplex Kit and Chip Kit (10× Genomics, V2 barcoding chemistry) and the subsequent steps were performed according to the manufacturer’s instructions. After the library preparation, sequencing reads were obtained by an Illumina NovaSeq 6000 sequencer. Cell Ranger (Zheng et al., 2017) was then used to map reads to human genome hg19 (Haeussler et al., 2019), and create count tables. After this, cells with less than 200 expressed genes, higher than 6,000 expressed genes, and/or mitochondria content higher than 5% were discarded. Lowly expressed genes [mean UMI (unique molecular identifiers per million reads) smaller than 0.1] were filtered out. Lastly, gene expression levels were normalized and log transformed for downstream analysis.

#### Homogeneity assessment

Kendall’s tau, a nonparametric measure of ordinal association (Hodges, JL Jr. and Lehmann, EL, 1963), was calculated between cell pairs within the sample to estimate the intrasample homogeneity. Specifically, 10,000 cell pairs within one sample were randomly selected, then Kendall’s tau of gene expression levels between cells in each pair was calculated. The average Kendall’s tau of 10,000 cell pairs was used to estimate the intrasample homogeneity. Samples with higher homogeneity are expected to have more similar expressional landscapes between cells, resulting in larger average values of Kendall’s tau.

#### Neuronal characteristics assessment

we utilized snRNA-seq data from postmortem brains (see details in Methods “Datasets used as references for the neuronal characteristics of ONCs” section) to predict the cell type of ONCs using *Seurat* (Stuart et al., 2019). Specifically, anchors between the reference (snRNA-seq data from postmortem brains) and query (scRNA-seq data from ONCs) datasets were identified and used for batch effect correction. Then the cell type of query cells was predicted based on the anchor weight, a measure of the shared neighbor overlap and distance between the anchor and query cells. As a quality control, we tested this analytic pipeline on samples with known cell types, including human brain neurons, human brain microglia, iPSC-derived neurons, and human PBMCs (see details in Methods “Datasets used as quality controls for *Seurat* cell type prediction” section), to ensure that correct predictions can be provided by this analysis.

### mRNA/protein expression and functional study of the glutamate receptor interacting protein 2 (GRIP2)

#### Quantitative reverse transcription-polymerase chain reaction (qRT-PCR)

We followed our published protocols (Sumitomo et al., 2018; Namkung et al., 2023). In brief, total RNA was isolated from the ONCs using the RNeasy Plus Mini Kit (Qiagen), and reverse-transcribed with a High-Capacity RNA-to-cDNA Kit (Applied Biosystems). All data were normalized with β-actin as a reference.

#### Western blotting

We followed our published protocols (Sumitomo et al., 2018; Takayanagi et al., 2021). Briefly, ONC whole cell lysates were subjected to SDS-PAGE followed by Western blotting with an anti-GRIP2 antibody (a kind gift from Dr. Richard L. Huganir, Johns Hopkins University) (Dong et al., 1999; Mejias et al., 2011) and an anti-alpha tubulin antibody (abcam, ab18251). Alpha tubulin levels were used as loading controls.

#### Surface and intracellular staining of AMPA-type ionotropic glutamate receptor / glutamate ionotropic receptor AMPA type subunit 2 (GRIA2)

We followed our published protocols (Namkung et al., 2023) with some modifications. GRIA2 tagged with a myc-epitope in the N-terminal extracellular region (a kind gift from Dr. Richard L. Huganir, Johns Hopkins University) (Anggono et al., 2013) were transfected to ONCs using Lipofectamine 3000, and ONCs were stained 2 days after transfection. To assess the surface myc-GRIA2, prior to cell permeabilization, ONCs were stained with a mouse monoclonal anti-myc antibody (Santa Cruz Biotech, sc-40). Then, ONCs were washed and fixed in 4% paraformaldehyde and incubated with a rabbit polyclonal anti-myc antibody (Santa Cruz Biotech, sc-789) in permeabilizing/blocking solution (0.5% Triton X, 4% normal goat serum, and 0.1% bovine serum albumin in PBS) to label intracellular myc-GRIA2. Alexa Fluor 488- and Alexa Fluor 568-conjugated secondary antibodies were used for the detection. Images were acquired on LSM700 confocal microscope (Zeiss) and analyzed using Image J software. The ratio of surface to total (surface + intracellular) fluorescence was calculated to measure the surface expression of GRIA2.

### Protein study of P62

We followed our published protocol for P62 Western blotting (Sumitomo et al., 2018). We used an anti-P62 antibody (MBL, PM066) and an anti-GAPDH antibody (Santa Cruz Biotech, sc-32233). GAPDH levels were used as loading controls.

### Statistical analysis

Analysis of covariance (ANCOVA) and partial correlation analysis were employed for group comparison and correlation analysis while controlling for confounding factors, respectively. Age, sex, race, and tobacco usage status (Yes/No) were controlled in all analyses. When only patients’ data were used, additional confounding factors including chlorpromazine equivalent dose and duration of illness were controlled.

## RESULTS

### ONCs are homogeneous

First, we asked whether ONCs are homogeneous. To address this question, we compared scRNA-seq data from ONCs of three healthy controls (see demographic summary in **Table 1**) with scRNA-seq data from immortalized cell lines (HEK293T and Jurkat cells) (Zheng et al., 2017), scRNA-seq data from iPSC-derived neural mixed culture (Earley et al., 2021), scRNA-seq data from iPSC-derived dopaminergic neurons (Earley et al., 2021), and snRNA-seq data from postmortem sACC and DLPFC from healthy subjects without neurological disorders (each individual subject was analyzed separately) (Tran et al., 2021). The immortalized cell lines were chosen as examples of homogeneous cells. Kendall’s tau, a measure of the similarity of expressional landscapes between cells within samples, was calculated to assess intrasample homogeneity.

**Table 1.**
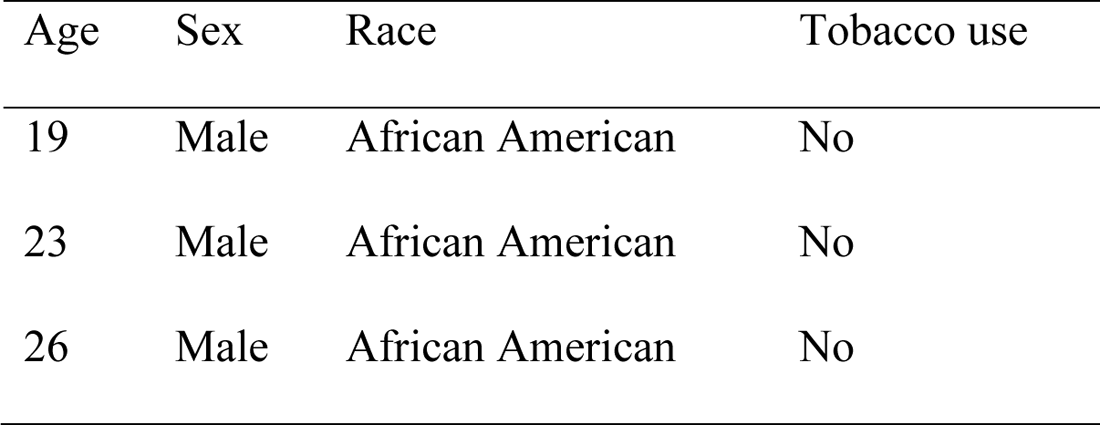
Demographic summary of healthy controls used for studying the homogeneous and neuronal characteristics of ONCs at the scRNA-seq data level.

| Age | Sex | Race | Tobacco use |
| --- | --- | --- | --- |
| 19 | Male | African American | No |
| 23 | Male | African American | No |
| 26 | Male | African American | No |

Our data showed that ONCs had similar homogeneity to well-established homogeneous cell lines such as HEK293T and Jurkat cells (**Fig. 2**). The average Kendall’s tau was 0.46, 0.44, and 0.42 for ONCs of each individual; similarly, Kendall’s tau was 0.44 and 0.46 for HEK293T and Jurkat cells, respectively. On the other hand, iPSC-derived neurons were more heterogeneous, with an average Kendall’s tau of 0.30, 0.36, 0.31, and 0.30 for iPSC-derived mixed neural culture, dopaminergic neurons, nociceptive sensory neurons, and spinal motor neurons, respectively. In the case of postmortem brain tissues with heterogeneous cell populations, the average Kendall’s tau was between 0.25 and 0.27. These results suggested that ONCs were close to homogeneous cell lines and more homogeneous than iPSC-derived neurons.

**Figure 2.**
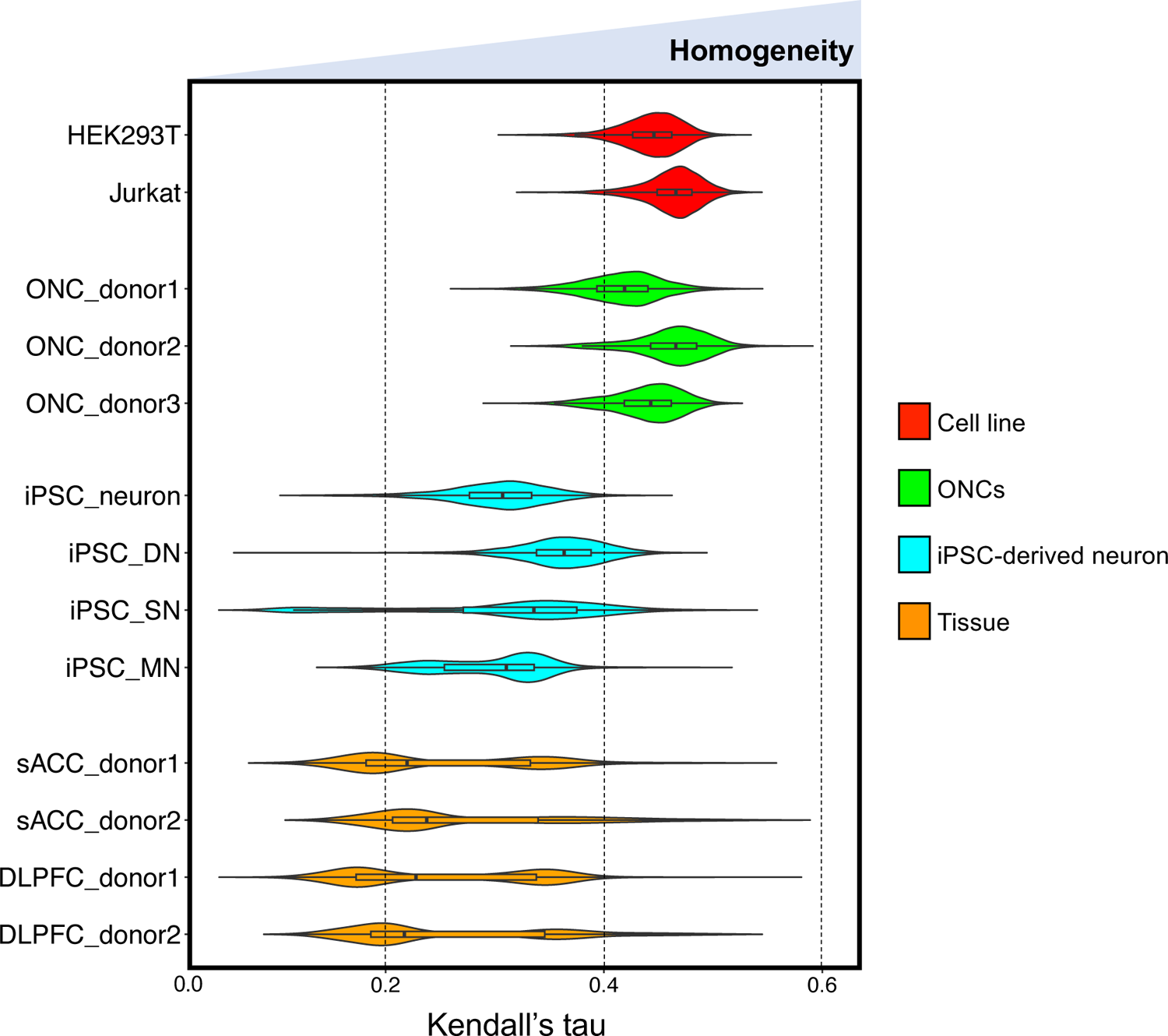
The homogeneity characteristics of ONCs. Kendall’s tau was calculated to assess the intrasample homogeneity. ONCs from three donors were compared with scRNA-seq data from homogeneous cell lines (HEK293T and Jurkat cells), scRNA-seq data from iPSC-derived cells including neural mixed culture (iPSC_neuron), dopaminergic neurons (iPSC_DN), nociceptive sensory neurons (iPSC_SN), and spinal motor neurons (iPSC_MN) and snRNA-seq data from postmortem brain tissues (individual subjects were analyzed separately: sACC_donor1, sACC_donor2, DLPFC_donor1, DLPFC_donor2)

### Neuronal characteristics of ONCs

Next, we asked whether the molecular signature of ONCs is neuronal. To address this question, we compared bulk RNA-seq and scRNA-seq data from ONCs with those from the brains, iPSC-derived dopaminergic neurons, and the OE tissues.

We conducted PCA analysis to compare bulk RNA-seq data from ONCs of 61 healthy controls (see demographic summary in **Table 2**) with bulk RNA-seq data from postmortem brains at different developmental stages, including the fetus, infancy, childhood, adolescence, and adulthood (Miller et al., 2014). Our analysis showed that ONCs were most similar to the fetal brain (Extended Data Fig. 3-1).

**Table 2.** Demographic summary of the cohort used for studying the neuronal characteristics of ONCs at the bulk RNA-seq data level. Abbreviations: SD, standard deviation; Y, yes; N, no; AA, African American.

|  | Controls (n=61) |
| --- | --- |
| Age (mean/SD) | 23.28 / 3.17 |
| Sex (Male/Female) | 31 / 30 |
| Race (AA/Caucasian/Asian/others) | 39 / 16 / 2 / 4 |
| Tobacco use (Y/N) | 2 / 59 |

We also utilized scRNA-seq data from ONCs of 3 healthy controls (**Table 1**) and snRNA-seq data from three independent postmortem brain datasets as references. *Seurat* predicted ONCs as neurons against other cell types in the brain, including astrocytes, microglia, oligodendrocytes, oligodendrocyte progenitor cells, and endothelial cells, for all three reference datasets (**Fig. 3A, 3B**; Extended Data Fig. 3-2).

**Figure 3.**
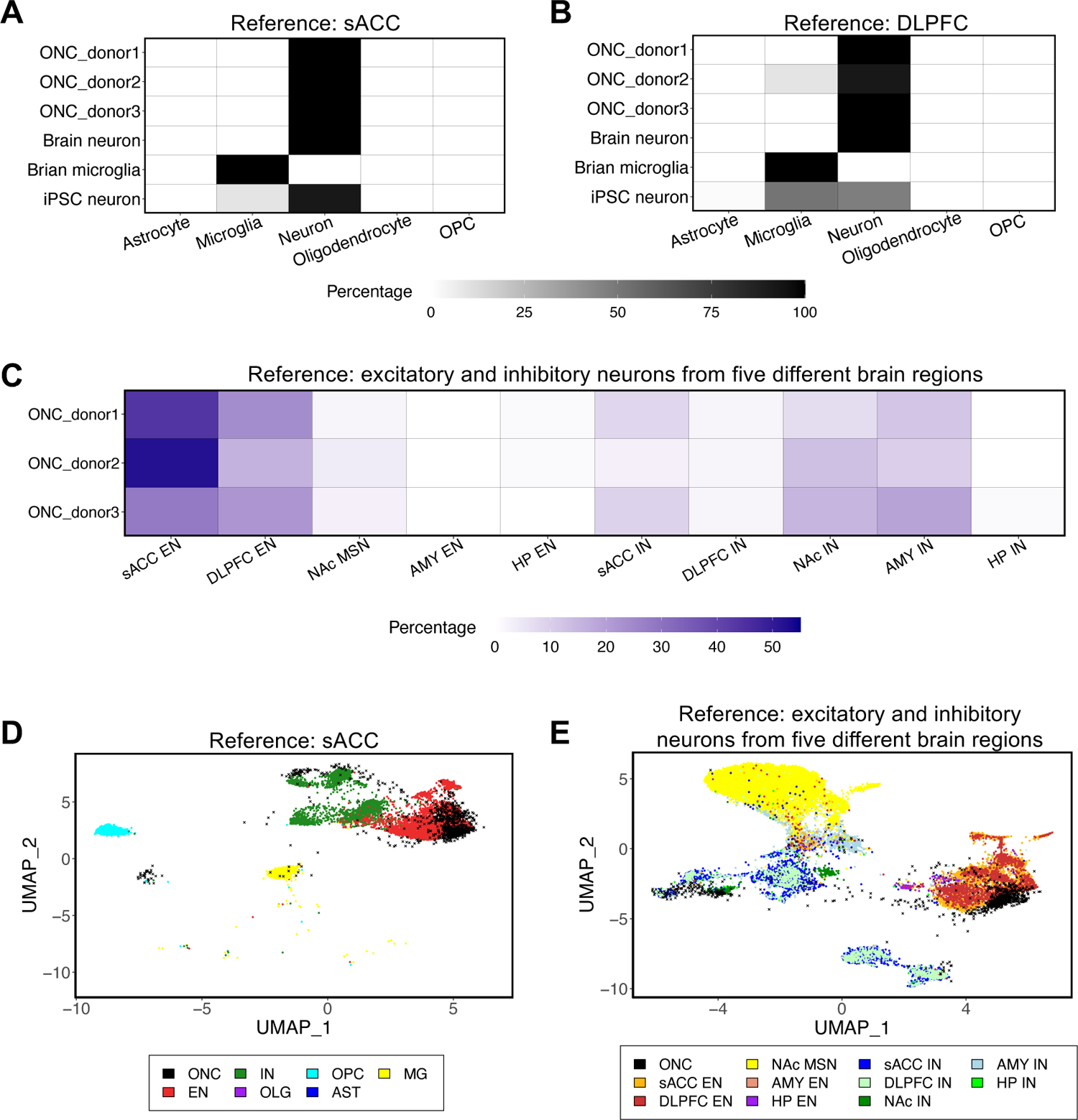
The neuronal characteristics of ONCs. (A) *Seurat* cell type prediction of ONCs against human brain cells in the sACC. Publicly available data from brain neurons, iPSC-derived dopaminergic neurons, and brain microglia were used for quality control. The heatmap shows the percentage of cells in the tested samples (y-axis) against the reference cell types (x-axis). (B) *Seurat* cell type prediction of ONCs against human brain cells in the DLPFC. (C) *Seurat* cell type prediction of ONCs against excitatory (EN) and inhibitory (IN) neurons across five different brain regions: sACC, DLPFC, hippocampus (HP), nucleus accumbens (NAc), and amygdala (AMY). (D) UMAP projection of ONCs on the 2D space of human brain cells in the sACC. (E) UMAP projection of ONCs on the 2D space of neuron subtypes in five brain regions.

When comparing ONCs with neuronal subtypes from five different brain regions, including the sACC, DLPFC, hippocampus, nucleus accumbens, and amygdala (Tran et al., 2021), *Seurat* predicted the majority of ONCs as excitatory neurons in the sACC and DLPFC (**Fig. 3C**).

UMAP (Uniform Manifold Approximation and Projection) (McInnes et al., 2018) also placed ONCs onto the space overlapped with excitatory neurons in the sACC and DLPFC (**Fig. 3D, 3E**).

Lastly, we compared scRNA-seq data from ONCs with those from the OE tissues (Durante et al., 2020). UMAP showed that ONCs were clustered in the space where horizontal basal cells gathered and far away from where pericytes and fibroblasts gathered (Extended Data Fig. 3-3).

Olfactory horizontal basal cells are stem cells that can regenerate olfactory neurons and non-neuronal cells after injuries (Iwai et al., 2008). Interestingly, despite their origin from the OE, ONCs resemble neurons in the brain (**Fig. 3**) rather than olfactory sensory neurons in the OE (Extended Data Fig. 3-3).

Together, our study showed that ONCs are homogeneous cells whose molecular signatures are similar to neuronal cells, which may be a useful cell resource for studying molecular signatures of brain disorders. Homogeneous cells are advantageous in research where bulk cells are needed, such as in protein studies.

### Unique utility of ONCs in a cell-based mechanistic study for the pathophysiology of TR in psychotic patients

A significant advantage of having homogenous cell cultures is that we could conduct cell-based mechanistic studies where single-cell resolution techniques are still limited. To obtain such a proof-of-concept, by using ONCs from patients, we focused on a molecular and cellular mechanism that might be specifically associated with TR in psychosis.

TR is the most critical medical question in psychotic disorders, resulting in a significant social burden (Kennedy et al., 2014; Nucifora et al., 2018). One-third of patients with psychosis fall into TR, leading to more devastating outcomes in the disease trajectory (Lally et al., 2016; Wimberley et al., 2016; Nucifora et al., 2018). Several groups have addressed brain imaging signatures associated with TR (Palaniyappan et al., 2013; Egerton et al., 2018; Dempster et al., 2020; Li et al., 2020; Kraguljac et al., 2021; Yang et al., 2022b). Nevertheless, the biological mechanism(s) underlying TR remains elusive. According to the criterion utilized in many published studies (see **Methods**), we stratified 62 patients whose ONCs were available into TR and non-TR groups (see demographic summary in **Table 3**).

**Table 3.** Demographic summary of the cohort used for studying the molecular mechanisms of TR in psychosis. Abbreviations: SD, standard deviation; Y, yes; N, no; AA, African American.

|  | TR (n=16) | non-TR (n=46) | p-value |
| --- | --- | --- | --- |
| Age (mean/SD) | 21.81 / 2.32 | 23.22 / 4.21 | 0.10 |
| Sex (Male/Female) | 12 / 4 | 33 / 13 | 1.00 |
| Race (AA/Caucasian/Asian/others) | 9 / 5 / 1 / 1 | 29 / 13 / 1 / 3 | 0.86 |
| Tobacco use (Y/N) | 8 / 8 | 7 / 39 | 0.01 |
| DOI (months, mean/SD) | 20.06 / 15.99 | 16.87 / 13.34 | 0.48 |
| CPZ dose (mg, mean/SD) | 367.64 / 268.12 | 268.32 / 215.70 | 0.23 |

First, we analyzed bulk RNA-seq data of ONCs from 16 TR and 46 non-TR patients, looking for differentially expressed genes (DEGs) associated with TR. After controlling confounding factors and multiple comparison correction, we identified seven DEGs (adjusted p-value < 0.05), where the DEG with the largest fold change was the gene encoding GRIP2 (**Fig. 4A**). GRIP2 is one of the GRIP family molecules that interact with AMPA-type ionotropic glutamate receptors, called GRIA proteins. The GRIP2 gene expression was reportedly decreased in the postmortem superior temporal gyrus in patients with schizophrenia (Bowden et al., 2008). The direction of this pathological change is consistent between ONCs and postmortem brains. Further enrichment analysis of top genes (p-value < 0.05 between TR and non-TR groups) showed that GWAS risk genes for schizophrenia and psychosis (see details in **Methods**) were significantly overrepresented in these genes (p-value = 2.66E-7). We also conducted a pathway analysis and identified 12 significant pathways between TR and non-TR patients, which included immune/inflammatory pathways (**Table 4**). These pathways have also been highlighted when we compared the entire patients (including both TR and non-TR patients) and healthy subjects in our past publications (Yang et al., 2021, 2022a).

**Figure 4.**
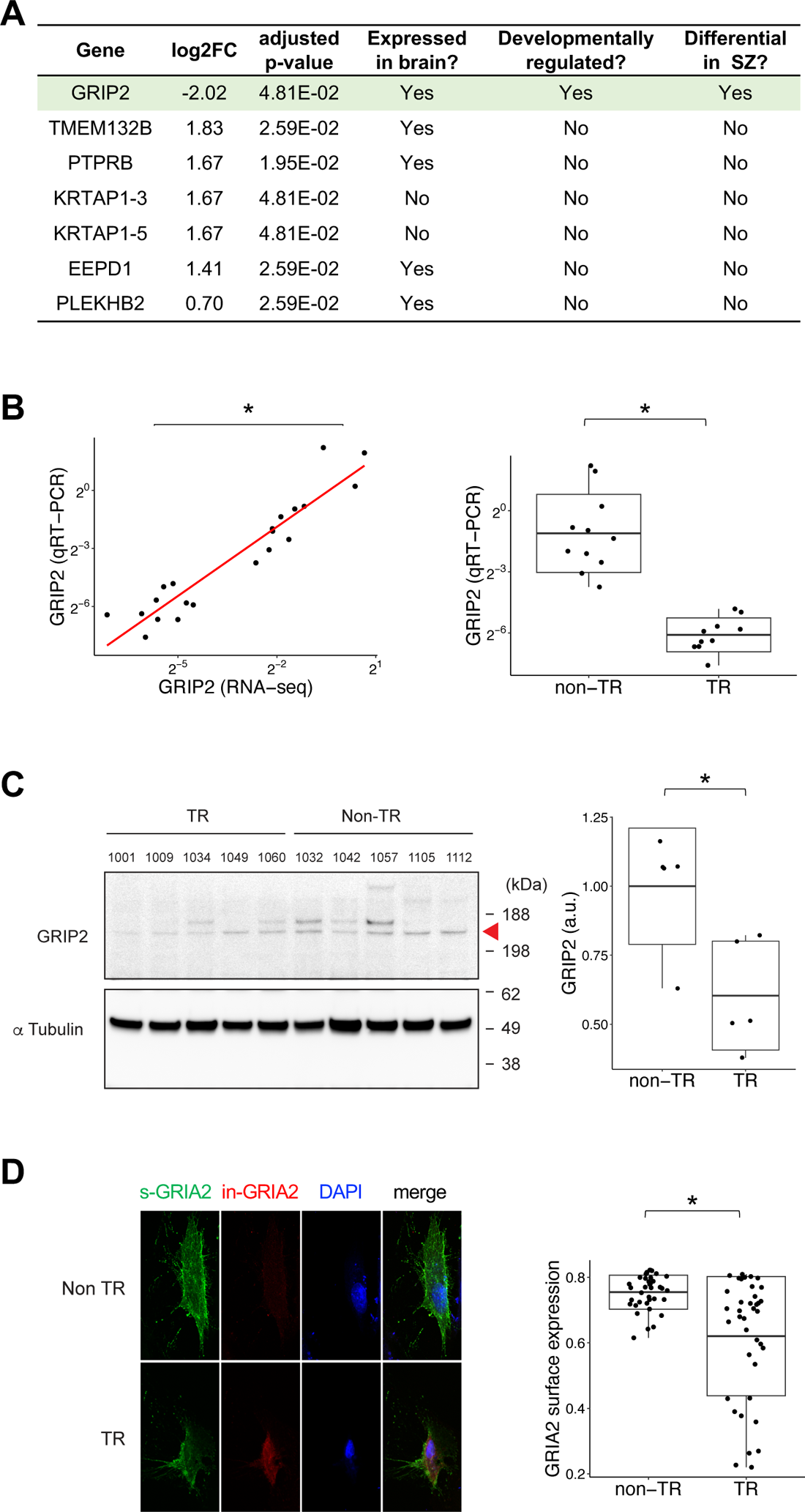
GRIP2 and TR in psychosis. (A) DEGs between TR (n=16) and non-TR (n=46) patients in ONC bulk RNA-seq data. (B) Confirmation of a significant reduction of GRIP2 expression in TR by qRT-PCR of a subset of patients (10 TR and 11 non-TR). The data between qRT-PCR and RNA-seq are significantly correlated. (C) Confirmation of a significant reduction of GRIP2 expression in TR by Western blotting of a subset of patients (5 TR and 5 non-TR). (D) A significant reduction of cell surface presentation of exogenously expressed myc-GRIA2 in ONCs from TR patients (n=3, total 38 replicates) compared with that in ONCs from non-TR patients (n=3, total 38 replicates). Symbol * denotes significant results (p-value < 0.05).

**Table 4.** Significant pathways altered in TR patients compared with non-TR patients. GSEA identified 12 significant pathways (adjusted p-value < 0.05). Abbreviations: NES, normalized enrichment score.

| Pathway | Catalog | NES | adjusted p-value |
| --- | --- | --- | --- |
| Ubiquitin-dependent degradation of cyclin D | Cell cycle | 1.97 | 3.68E-02 |
| XBP1(S) activates chaperone genes | Cellular responses to stimuli | 2.40 | 0.00E+00 |
| Hh mutants are degraded by ERAD | Disease | 2.06 | 1.96E-02 |
| Defective CFTR causes cystic fibrosis | Disease | 2.00 | 2.78E-02 |
| Interleukin-35 signaling | Immune system | 2.11 | 2.08E-02 |
| Interleukin-10 signaling | Immune system | 2.08 | 1.83E-02 |
| Interleukin-4 and interleukin-13 signaling | Immune system | 2.02 | 2.52E-02 |
| Cross-presentation of soluble exogenous antigens endosomes | Immune system | 1.94 | 4.75E-02 |
| Cholesterol biosynthesis | Metabolism | 1.94 | 4.53E-02 |
| Hedgehog ligand biogenesis | Signal transduction | 2.10 | 1.63E-02 |
| COPI-mediated anterograde transport | Vesicle-mediated transport | 2.05 | 2.06E-02 |
| COPI-dependent Golgi-to-ER retrograde traffic | Vesicle-mediated transport | 1.91 | 4.98E-02 |

We next conducted qRT-PCR to validate the decrease of GRIP2 expression in ONC bulk RNA-seq data. We observed a significant correlation between the qRT-PCR and bulk RNA-seq levels (p-value = 2.17E-5) (**Fig. 4B**). Consistently, we also observed a significant decrease (p-value = 4.08E-5) in GRIP2 in TR patients compared with non-TR patients through qRT-PCR (**Fig. 4B**). Furthermore, we conducted Western blotting and confirmed this TR-associated decrease in GRIP2 at the protein level (p-value = 0.02) (**Fig. 4C**).

Based on the above findings, we further hypothesized that the significant reduction of GRIP2 protein might affect the trafficking of AMPA receptors (GRIA proteins) in TR patients. The functional impact of GRIP proteins on their protein interactor GRIA2 has been studied in cell cultures, in which an alteration of cell surface GRIA2 is used as a key outcome measure (Osten et al., 2000; Kulangara et al., 2007). Thus, to validate this hypothesis, we measured the cell surface expression of GRIA2 and observed a significant decrease in the surface expression of GRIA2 in TR patients compared with non-TR patients (p-value = 3.53E-4) (**Fig. 4D**).

Taken together, our study showed a reduction of GRIP2 expression and a resultant change of GRIA2 in ONCs from TR patients, in comparison with those from non-TR patients. These support the utility of patient ONCs to explore and validate a clinical feature-associated biological mechanism.

### Unique utility of ONCs in a protein study for the pathophysiology of aging in psychotic patients

Homogenous cell cultures, such as ONCs, are useful for protein assays in which single-cell-based approaches are still limited. Previously, by utilizing ONCs, we observed increased levels of P62 protein in patients with schizophrenia compared with those from healthy controls (Sumitomo et al., 2018). P62 is a hub protein that plays a role in autophagy-related cellular processes (Tomoda et al., 2020). When the process is attenuated, the P62 protein level increases (Tomoda et al., 2020).

In the present study, by using a new cohort with an increased number of subjects (33 healthy controls and 30 psychotic patients, see **Table 5**) and ANCOVA controlling for age, sex, race, and tobacco usage status (Yes/No), we observed an upregulation of P62 protein in ONCs from patients with psychotic disorders compared with healthy controls (p-value = 3.53E-3) (**Fig. 5A**). This observation was consistent with our past publication (Sumitomo et al., 2018), confirming the original observation in a different cohort. Thus, we conducted further examinations of P62 protein in association with schizophrenia.

**Figure 5.**
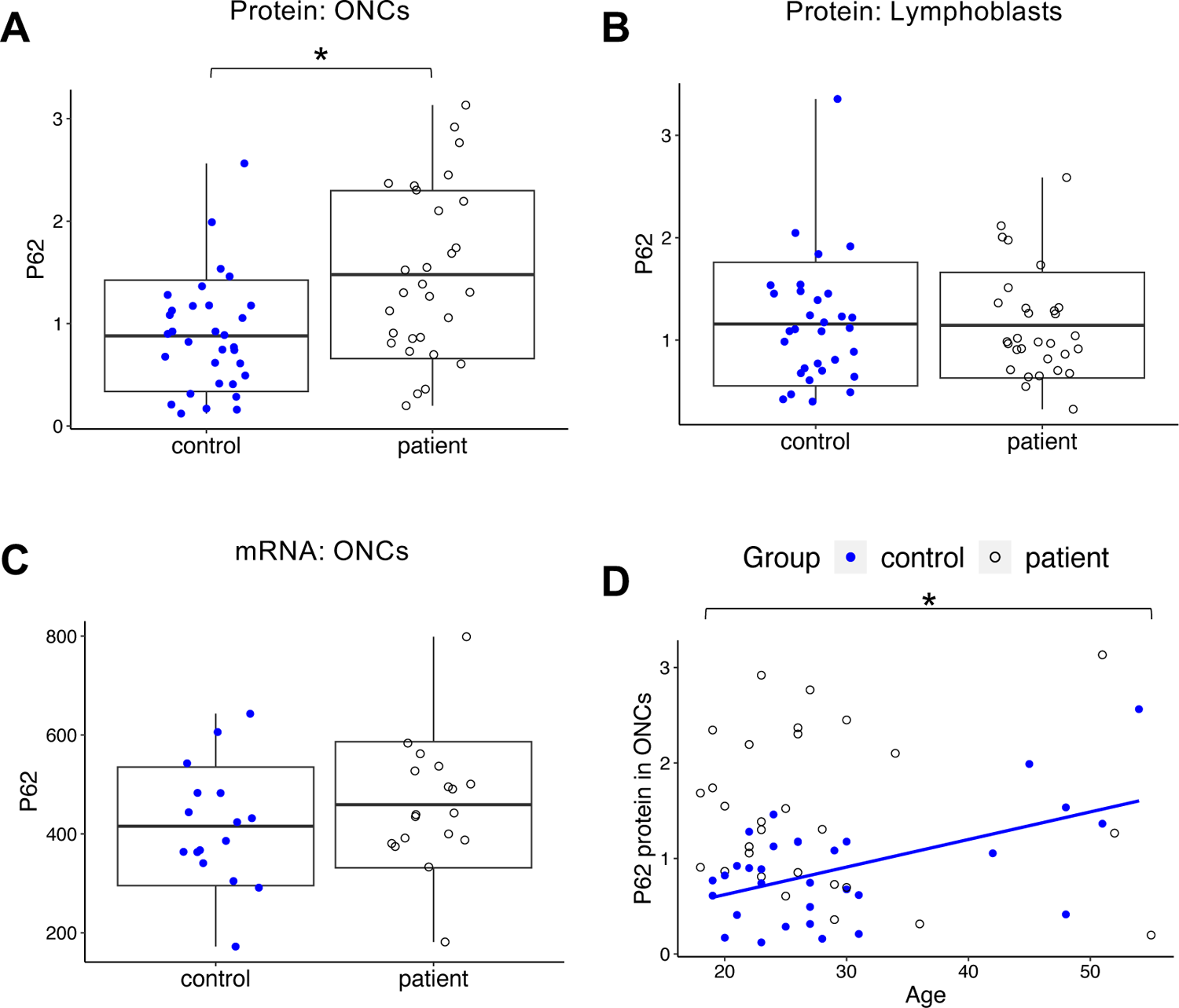
P62 and psychotic disorders. (A) A significant increase of P62 protein levels in ONCs in psychotic patients (n=30) compared with healthy controls (n=33). (B) No significant difference in P62 protein level in lymphoblasts between psychotic patients (n=30) compared with healthy controls (n=32, one control didn’t donate lymphoblasts) (C) No significant difference in P62 mRNA levels in ONCs between psychotic patients (n=18, we didn’t have mRNA data from 12 patients) and healthy controls (n=16, we didn’t have mRNA data from 17 controls) (D) A significant correlation between age and P62 protein levels in ONCs in healthy controls (n=33); no significant correlation in psychotic patients (n=30). Symbol * denotes significant results (p-value < 0.05).

**Table 5.** Demographic summary of the cohort used for studying P62 levels. Abbreviations: SD, standard deviation; Y, yes; N, no; AA, African American.

|  | Controls (n=33) | Patients (n=30) | p-value |
| --- | --- | --- | --- |
| Age (mean/SD) | 28.79 / 9.93 | 27.90 / 9.71 | 0.72 |
| Sex (Male/Female) | 22 / 11 | 20 / 10 | 1.00 |
| Race (AA/Caucasian/Asian/others) | 21 / 10 / 1 / 1 | 21 / 8 / 1 / 0 | 0.78 |
| Tobacco use (Y/N) | 7 / 26 | 16 / 14 | 0.02 |

This disease-associated protein change may be unique to neuronal cells, as we did not observe an upregulation of P62 protein in lymphoblasts from the same set of patients (p-value = 0.94) (**Fig. 5B**) We did not observe an upregulation of P62 at the mRNA level in the same set of ONCs (p-value = 0.53) (**Fig. 5C**), which is consistent with the notion of P62 regulation at the posttranscriptional levels (Klionsky et al., 2012). Given that the autophagic process is attenuated with aging, we examined whether the levels of P62 in ONCs may be affected by aging via partial correlation analysis controlling for sex, race, and tobacco usage status. We observed a significant age-dependent increase in P62 protein in ONCs from healthy subjects (p-value = 4.63E-3) (**Fig. 5D**). Interestingly, when we examined ONCs from patients with psychotic disorders, such an age-dependent phenotype was not observed (p-value = 0.84) (**Fig. 5D**). The levels of P62 were already high in young adulthood, and no further upregulation was observed in older patients.

The present study on P62 protein shows the reproducibility and utility of ONCs in biochemical protein studies that require cell homogeneity. Additionally, our findings suggest that ONCs have the potential to uncover disease-associated neuronal mechanisms.

## DISCUSSION

In the present study, we introduced a unique and easy-to-perform protocol that could establish neuronal cells from the OE. Systematic analysis using single-cell and bulk RNA-seq data showed that our culture protocol could produce homogeneous neuronal cells, which also provided strong support for many past studies that used ONCs (Sawa and Cascella, 2009; Horiuchi et al., 2013; Kano et al., 2013; Mor et al., 2013; Lavoie et al., 2017; Sumitomo et al., 2018; Takayanagi et al., 2021; Namkung et al., 2023). We propose that ONCs can be used as neuronal surrogates to address brain functions and disorders, particularly in pharmacological and biochemical studies in which the homogeneity of cells is crucial for assays.

ONCs have several additional advantages. As these are biopsied from living subjects, ONCs allow for studies linking neuronal molecular profiles to clinical manifestations (Jaaro-Peled et al., 2022; Mihaljevic et al., 2022). As the olfactory neural epithelium can regenerate quickly, repeated biopsies are possible. In more than one institution, studies with repeated biopsies have been approved and published (Sattler et al., 2011; McLean et al., 2018). Repeated biopsies are particularly advantageous in clinical trials when assessing the treatment response of target molecules in neuronal cells. We can also examine the molecular changes in correlation with clinical amelioration by conducting biopsies before and after trials: for example, symptom changes and transcriptional regulation of Collapsin Response Mediator Protein 1 (CRMP1) in olfactory neuronal cells have been reported in patients with bipolar disorder in response to lithium treatment (Sattler et al., 2011; McLean et al., 2018). Furthermore, ONCs obtained from repeated biopsies in a longitudinal study could provide mechanistic insight associated with dynamic changes along the disease trajectory. In our psychosis cohort (Yang et al., 2022b; Wang et al., 2023), the return rate of study participants is equivalent between those who underwent nasal biopsy and those who did not. This indicates that the process of nasal biopsies is not invasive enough to discourage study participants in clinical studies. In addition, in theory, ONCs can be more easily scaled up for large populations with a lower budget and in less time when compared to iPSC-derived neurons. This scalability also enhances the translational potential of ONCs.

We also acknowledge that, compared to postmortem brain and iPSC-derived neurons, ONCs have several limitations. Although one group has published calcium signaling deficits in nasal biopsied olfactory primary neurons from patients with bipolar disorder (Hahn et al., 2005), ONCs and related cells do not show clear action potentials seen in mature neurons. ONCs are not suitable for studying the characteristics of mature and differentiated neurons. Recent advances in iPSC-based neurobiology have overcome these limitations.

In neurodegenerative disease research, postmortem brains have been used as a gold standard to explore molecular and cellular changes (Nagatsu and Sawada, 2007; Hopperton et al., 2018; Li et al., 2019). However, when we study neurodevelopmental disorders, we need to pay attention to at least two points: first, different from neurodegenerative diseases, pathophysiological processes may be active many years prior to an individual’s death, and may not remain after death. Second, in psychiatric disorders of neurodevelopmental origins, such as schizophrenia, long-term medication may modify intrinsic molecular signatures (Rapoport et al., 2012; Birnbaum and Weinberger, 2017). In addition, postmortem brains cannot be used for investigating disease trajectories where longitudinal studies are needed. To overcome these limitations associated with postmortem brain studies for neurodevelopmental disorders, ONCs through nasal biopsy in early active stages of the diseases may be particularly useful.

Accordingly, ONCs and related neuronal cells from nasal biopsy have been published more frequently in research for psychotic disorders, such as schizophrenia (Fan et al., 2012; Kano et al., 2013; Mor et al., 2013; Rhie et al., 2018; Sumitomo et al., 2018; Evgrafov et al., 2020; Takayanagi et al., 2021; Namkung et al., 2023). Although complete consensus has not been obtained yet, given the relatively small sample sizes of each past study, common pathways such as cell cycle control, calcium signaling, and immune/inflammatory pathways have been highlighted. One potential caveat is that the pathological changes in ONCs can reflect not only impairments underlying neuronal cells in general in the patients (the rationale for using ONCs as a surrogate of brain neurons) but also those accompanied by changes in the nasal cavity. For example, significant changes in the immune/inflammatory pathways in ONCs of psychotic patients (Yang et al., 2021) may be a converged outcome of intrinsic susceptibility to these pathways in patients with an additional local pathology elicited by air pollution and viral infection (risk factors for these diseases).

Nevertheless, in the present study, we demonstrated examples that could support the unique utility of ONCs as a neuronal resource obtained from living patients to directly study clinical feature-associated neuronal signatures and biological mechanisms. We proposed that a signaling cascade of the AMPA*-*type ionotropic glutamate receptor underlies the pathophysiology of TR in psychosis. Although TR is one of the most critical questions in psychosis studies and biological psychiatry in general, the shortage of neuron-relevant samples from living patients who indeed display TR in contrast to those from non-TR patients has slowed down the biological assessment for this question. ONCs can provide a breakthrough to this. ONCs may also be useful in studying the potential link of the OE impairment to resultant mental dysfunction elicited by SARS-CoV-2.

In summary, by conducting molecular characterization of ONCs using frontline next-generation sequencing techniques, we describe both advantages and limitations of ONCs, and propose their unique utility that can complement the use of iPSC-derived neurons and postmortem brains.

## Acknowledgments

this study is supported by National Institutes of Health Grants MH-092443 (to AS), MH-094268 (to AS), MH-105660 (to AS), MH-107730 (to AS), DC-016106 (to AL), AI-132590 (to AL), DA-041208 (to AK), AG-065168 (to AK), and MH-128765 (to AK); foundation grants from Stanley (to AS), RUSK/S-R (to AS), and a NARSAD young investigator award from Brain and Behavior Research Foundation (to AS, KY). The authors thank Dr. Sandra Lin for conducting the nasal biopsy.

