## Supplemental figures for "Human olfactory neuronal cells through nasal biopsy: molecular characterization and utility in brain science"

**Extended Data Figure 3-1. Comparison of bulk RNA-seq data between ONCs and postmortem brains at different developmental stages.**

PCA analysis was conducted and ONCs were projected onto the 2D space of postmortem brains at different developmental stages (fetus, infancy, childhood, adolescence, and adulthood) from BrainSpan.

**Extended Data Figure 3-2. Comparison of scRNA-seq data from ONCs with snRNA-seq data from human brain cells.**

(**A**) *Seurat* cell type prediction of ONCs against human brain cells in the sACC. Publicly available data from brain neurons, iPSC-derived dopaminergic neurons, brain microglia, and PBMC were used for quality control. The heatmap shows the percentage of cells in the tested samples (y-axis) against the reference cell types (x-axis).

(**B**) *Seurat* cell type prediction of ONCs against human brain cells in the DLPFC.

(**C**) *Seurat* cell type prediction of ONCs against human brain cells in the NAc.

(**D**) *Seurat* cell type prediction of ONCs against human brain cells in the HP.

(**E**) *Seurat* cell type prediction of ONCs against human brain cells in the AMY.

(**F**) *Seurat* cell type prediction of ONCs against human brain cells in the middle temporal gyrus (MTG).

(**G**) *Seurat* cell type prediction of ONCs against human brain cells in the prefrontal cortex (PFC)

**Extended Data Figure 3-3. Comparison of scRNA-seq data between ONCs and OE.**

UMAP was used to visualize the OE data in the 2D space. Both plots that include (right panel) and exclude (left panel) the projection of ONCs are shown here.

**Extended Data Figure 3-1.**

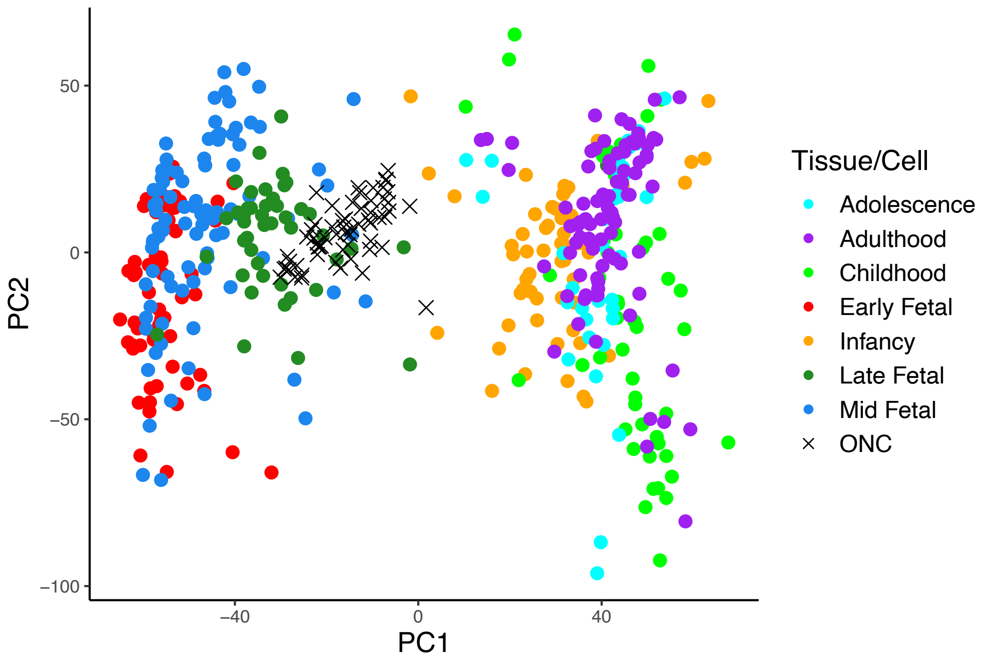

### Extended Data Figure 3-2.

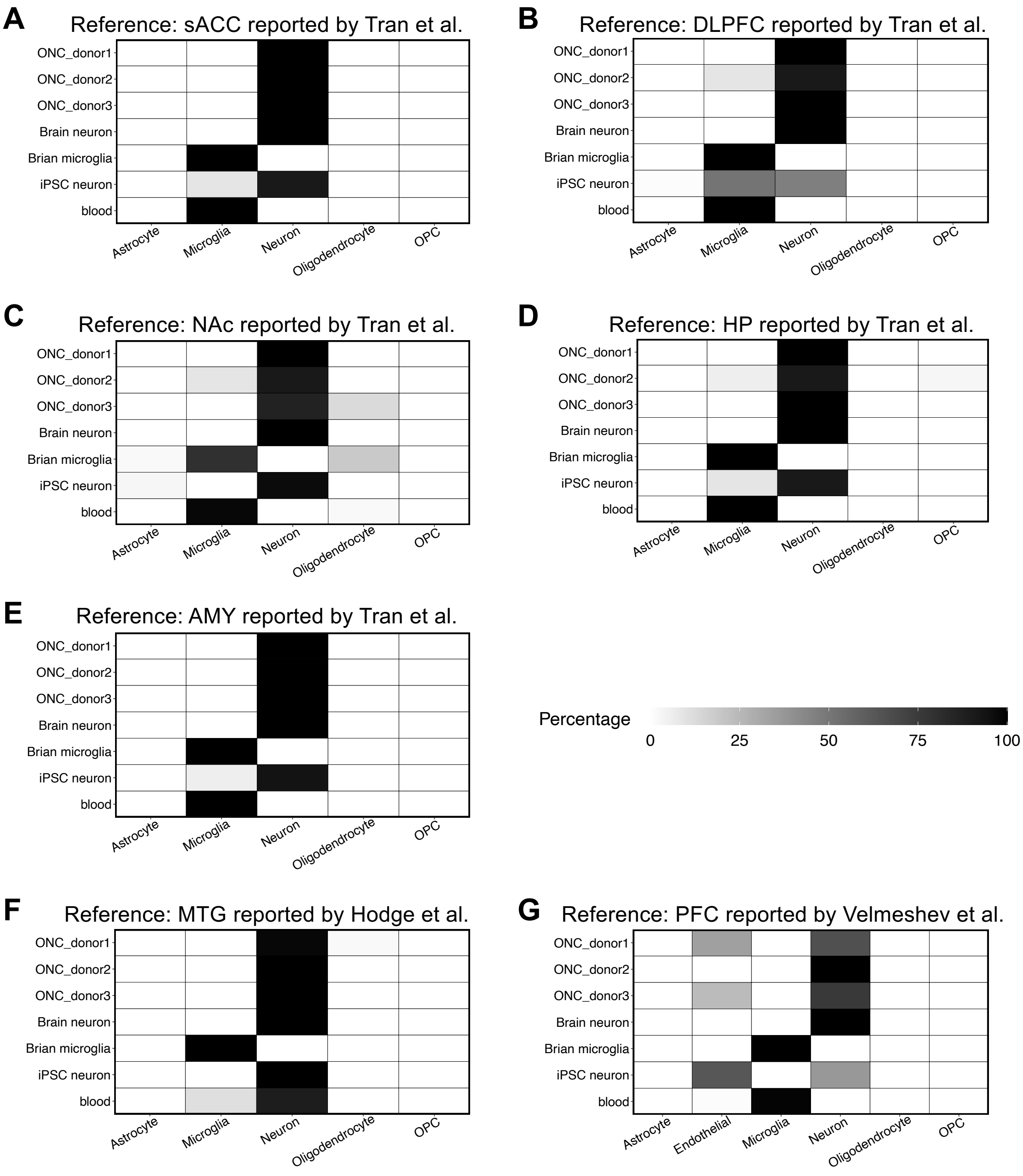

**Extended Data Figure 3-3.**

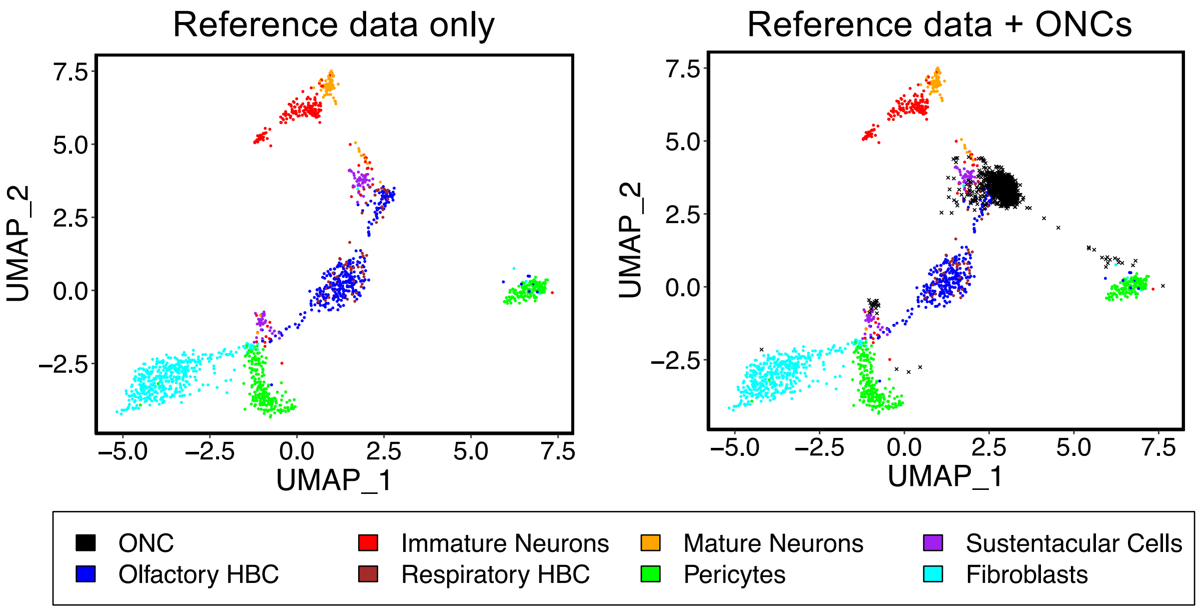
